# NanoMod: a computational tool to detect DNA modifications using Nanopore long-read sequencing data

**DOI:** 10.1101/277178

**Authors:** Qian Liu, Daniela C. Georgieva, Dieter Egli, Kai Wang

## Abstract

**Background:** Recent advances in single-molecule sequencing techniques, such as Nanopore sequencing, improved read length, increased sequencing throughput, and enabled direct detection of DNA modifications through the analysis of raw signals. These DNA modifications include naturally occurring modifications such as DNA methylations, as well as modifications that are introduced by DNA damage or through synthetic modifications to one of the four standard nucleotides.

**Methods:** To improve the performance of detecting DNA modifications, especially synthetically introduced modifications, we developed a novel computational tool called NanoMod. NanoMod takes raw signal data on a pair of DNA samples with and without modified bases, extracts signal intensities, performs base error correction based on a reference sequence, and then identifies bases with modifications by comparing the distribution of raw signals between two samples, while taking into account of the effects of neighboring bases on modified bases (“neighborhood effects”).

**Results:** We evaluated NanoMod on simulation data sets, based on different types of modifications and different magnitudes of neighborhood effects, and found that NanoMod outperformed other methods in identifying known modified bases. Additionally, we demonstrated superior performance of NanoMod on an *E. coli* data set with 5mC (5-methylcytosine) modifications.

**Conclusions:** In summary, NanoMod is a flexible tool to detect DNA modifications with single-base resolution from raw signals in Nanopore sequencing, and will greatly facilitate large-scale functional genomics experiments in the future that use modified nucleotides.

## Background

An important type of covalent modification in epigenetics is DNA modification, where a chemical residue can be added to one of the four standard nucleotides (A, C, G, T) in a DNA molecule [1]. Those added residues can be methyl, carboxyl, ethyl, formyl, hydroxymethyl, dimethyl groups and other larger chemicals such as biotin and Idoxuridine, resulting in various types of DNA modifications. The DNA modifications can exist naturally in genomes or can be introduced synthetically into DNA molecules for research purposes. For example, DNA methylation, a common and well-studied type of modification, is formed when a methyl group is added into the adenines or cytosines in a DNA molecule, and different types of methylations exist depending on which atomic position in an adenine or cytosine is modified, such as 5-methylcytosine (5mC) and N6-methyladenosine (6mA). Various naturally occurring DNA modifications have been widely discovered in all kingdoms of life [2]. They play a critical role in regulating cellular states and functions, controlling which genes are turned on/off, dramatically affecting gene expression and eventual production of proteins and their functions [3]. In comparison, synthetically introduced DNA modifications can mark specific positions in genome sequence, facilitating functional genomics studies. For example, labeling specific DNA sequence motifs by fluorescence signals in a genome can facilitate optical mapping of genomes and the detection of structural variants [4]. Furthermore, incorporation of modified DNA bases during DNA synthesis can be used to track patterns of DNA replication in a genome-wide scale through optical mapping [5]. However, there are currently no genome-wide methods that allow the detection of replicated and non-replicated DNA with base-pair resolution.

Several different genomic techniques have been developed to detect DNA modifications, especially for DNA methylations. For example, bisulfite sequencing is a widely used method for detecting DNA methylations, where unmethylated cytosines are converted to uracil and Illumina short-read sequencing techniques are used to call methylated and unmethylated cytosines from sequence data [6]. However, the harsh process in bisulfite treatment results in a large fraction of DNA fragmentation, which thus requires large quantity of DNA and complicated the analysis of highly variable, heterogeneous epigenome [3]. Immunoprecipitation together with Illumina short-read sequencing were also used to detect DNA or RNA modifications [7, 8], but these methods can detect only broad genomic regions with methylation without single base resolution. Other studies took advantage of PacBio single-molecule real-time (SMRT) sequencing techniques to directly detect DNA modifications using the principle that the existence of DNA modifications would affect DNA polymerase kinetics during SMRT sequencing [9-12]. Modifications in RNA can also be detected using PacBio SMRT sequencing [13]. However, there was reduced signal-to-noise ratio for 5mC modifications[14] and the improved enzymatic treatment of 5mC detection using Tet1[15] also had incomplete and context-dependent treatment[3]. A comprehensive review can be found in [16].

Recent studies have explored the use of Oxford Nanopore sequencing techniques for the detection of DNA modifications. In Nanopore sequencing, electric current change occurs when a k-mer passes through a nanopore, and different molecules (such as standard nucleotides and their modified versions) generate different current change, depending on sequence contexts. Several prior studies [17, 18] have carefully analyzed ionic current signals and demonstrated the feasibility of using Nanopore signals to identify DNA modifications by comparing current levels of methylated (that is, 5mC and 5- hydroxymethylcytosine (5hmC)) DNA copies with current levels of unmethylated DNA copies. They found that more C5-cytosine variants (1 unmethylated cytosine and 4 cytosine modifications) could also be identified using Nanopore sequencing data with higher accuracy in a background of known sequences [19]. Recently, three groups have quantified the strength of using Nanopore platform for detecting DNA modifications at a large scale [3, 20, 21]: Simpson *et. al.* developed a HMM (hidden Markov model) to distinguish 5mC from cytosine[3] in *E. coli* and *Homo sapiens* and integrated it in nanopolish, but this method cannot detect non-CpG methylations; Mclntyre *et.al.* designed mCaller to improve the detection of 6mA and tested the 6mA detection in mouse, *E. coli* and Lambda phage DNA [20]; Rand *et. al.* analyzed three types of cytosine (*i.e.*, cytosine, 5mC and 5hmC) and also 6mA in *E. coli* with different phases using HMM with a hierarchical Dirichlet process, with an implementation in the signalAlign package [21]. The results demonstrated feasibility to achieve improved performance in detecting DNA modifications [3, 20, 21], but they needed large prior training datasets for HMM [2], and therefore cannot be extended for detecting different types of modifications (especially synthetically introduced modifications). Stoiber *et. al.* proposed MoD-seq in the nanoraw package to identify modifications in the absence of large prior training dataset [2]. Here we developed NanoMod to achieve improved performance in the detection of modified bases even in the absence of any training data, though NanoMod can optionally leverage existing training data to further improve performance.

NanoMod was designed for the detection of *de novo* DNA modifications (for example, synthetically introduced modifications). The inputs of NanoMod were a group of reads from a DNA sample with modification at specific bases and a group of reads from the matched non-modified sample. The nucleotide sequence for the sample is assumed to be known, that is, the reference genome must be already known *a priori*. Currently, within NanoMod, we used albacore for basecalling, and then perform an indel error correction by aligning the events of electric signals to a reference genome, similar to the procedure implemented in nanoraw. After that, two groups of electric signals for each genomic position were compared using the Kolmogorov-Smirnov test [22] in a per-base level to identify bases with significantly different distributions of signals between the two groups. Finally, weighted Stouffer’s method was used to combine the effects of neighboring bases since some modifications (especially bulky ones) may have strong neighbor effects that affect electric signals in neighboring non-modified bases. We evaluated NanoMod on simulation data of modifications with different properties and on a published *E. coli* methylation data set.

## Methods

### NanoMod

The input of NanoMod is a dataset with two groups of reads: one from a sample with DNA modifications at specific positions and the other is the matched non-modified sample. The output is the ranked list of positions with potential modifications, as shown in **Figure 1**. NanoMod does not require prior training data, but it cannot detect the specific type of modification either. However, given a large-scale data set with known modifications at known positions, it is possible to use them as prior information to train a model and analyze a new dataset with the same type of modifications by NanoMod. The several steps involved in NanoMod are illustrated below.

**Figure 1.**
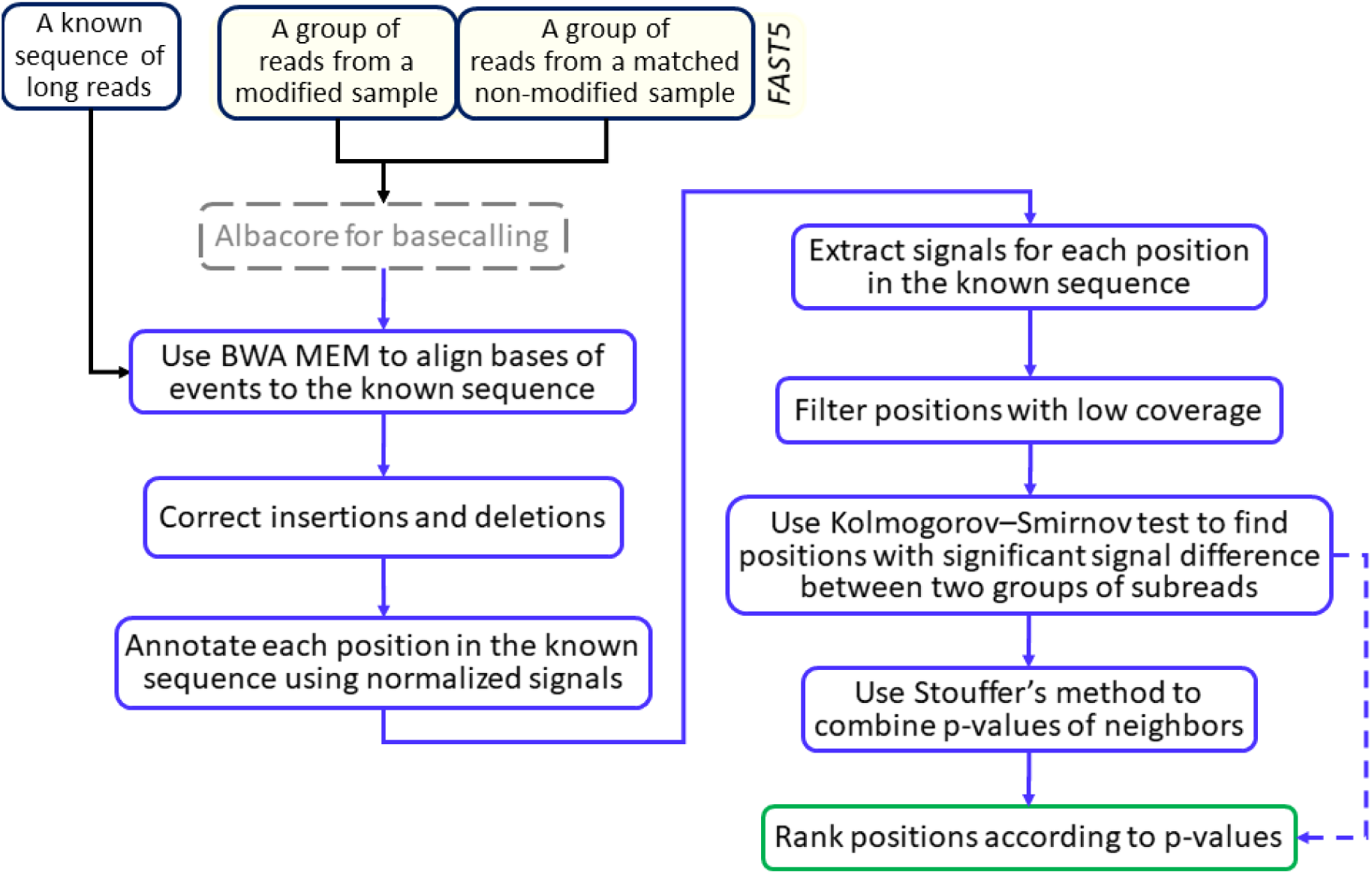
The flowchart of NanoMod. The squares with dotted line refer to components that require external tools, while the dotted arrow line suggests an alternative solution.

### Basecalling by albacore

Nanopore raw data on a long read consists of a time series of raw signals measured by the Oxford Nanopore sequencer such as MinION or GridION. Each raw signal is a digital integer value, a measure of the changes of electric current when a k-mer (for example, 5-mer) passes through nanopores. Since the acquisition frequency is usually much higher than the speed of translocation of bases passing through nanopores, the same k-mer may be measured multiple times when it passes through the pore. Since the speed of translocation is not constant, different k-mers may have different numbers of measurements. More importantly, errors and noises may exist during signals acquisition on the k-mers, making the precise interpretation of bases from raw signals more challenging. In other words, given a set of electric signals when a DNA molecule passes through the pore, it is not straightforward to convert them directly into a series of nucleotides.

To generate bases from Nanopore signals, raw signals are typically segmented into separate “events” in albacore (Note that the latest version albacore uses raw signals for basecalling, thus the segmentation step is no longer needed.) Each event consists of a consecutive series of raw signals that significantly deviate from the two direct neighboring events. The joint analysis of neighboring events with multiple overlapping bases would finally generate a sequence of bases with the highest probability, which is a procedure that uses deep recurrent neural network as implemented in albacore. The output of albacore contains a read from a FAST5 file and the signal information of all its bases.

### Error correction and signal annotation

Long reads generated on the Nanopore platform usually have high error rates which may negatively affect downstream analysis. Since we assume that a reference genome is already available (*i.e.* the true nucleotide identity is assumed to be known in advance), to correct the base calling errors, BWA-MEM [23] was used to align Nanopore long reads to the known sequence, and then the indels (possible basecalling errors) were corrected by a re-segmentation process which is similar to the indel correction procedure in nanoraw[2]. An insertion error suggests that two adjacent segmented events might be from the same k-mer, and thus, one of the two neighbor events of the insertion is merged with the insertion event for generating a new neighbor event. A deletion error suggests that the neighboring events of the deletion are be generated by one additional k-mer, and thus, the several closest neighbor events of the deletion are re-segmented so that one additional event can be generated. When the neighboring events to be re-segmented contain other indels, the collection of events are first merged together and then re-segmented so that proper events can be generated. The number of neighboring events is automatically determined so that there are enough number of signal measurements for each event after the re-segmentation. Meanwhile, to address the issue of homopolymer error, if there are *L*_*r*_ > 5 single nucleotide repeats in the sequence, the middle *L*_*r*_ − 4 new positions would share a certain new event after re-segmentation.

To illustrate this further, examples of the deletion correction procedure and insertion correction procedure are shown in **Figure 2** and **Figure 3**, respectively. In **Figure 2**, there are two deletions. To generate the correct events in **Figure 2**, we grouped the two deletions together (shadowed region in green) with one upstream adjacent neighbor and one downstream adjacent neighbor. We then re-segmented those signals associated with the bases in the shadowed region, and obtain two additional events from the correction procedure. In **Figure 3**, we grouped the insertion event, one upstream adjacent neighbor and one downstream adjacent neighbor (shadowed region in yellow), and then re-segmented the signals to generate two events from the correction procedure.

**Figure 2.**
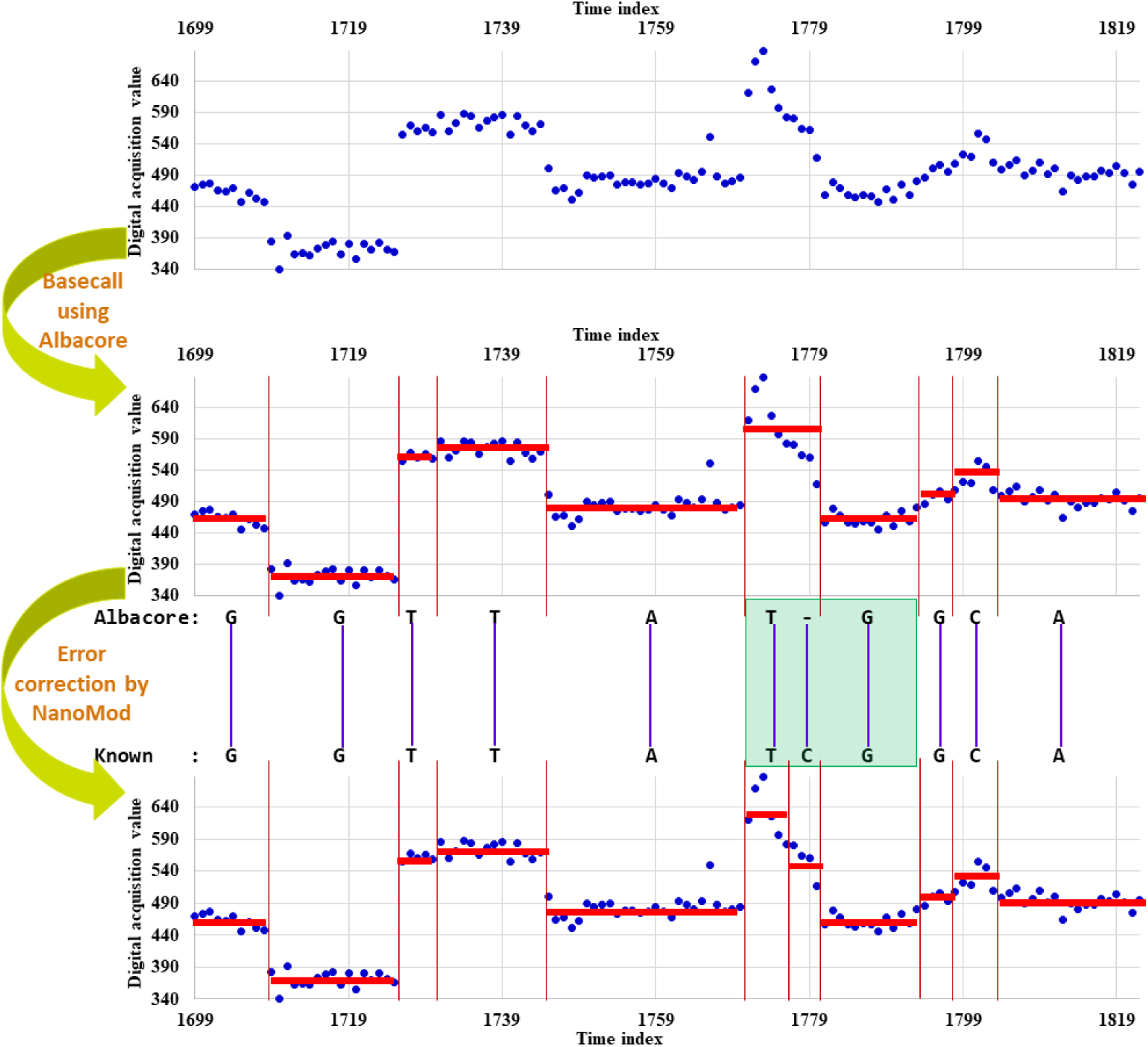
*An example of the deletion correction procedure in NanoMod*. *X axis represents time of signal acquisition, and y axis denotes detected signal values by Nanopore sequencers before standardization. ‘Albacore’ represents a sequence of bases called based on original events before error correction, and ‘Known’ represents the known sequence. Each red horizontal bar represents an event split by vertical lines. ‘-’ in ‘Albacore’ suggests a deletion. The region shadowed in green shows the deleted bases together with one upstream and one downstream neighbors.*

**Figure 3.**
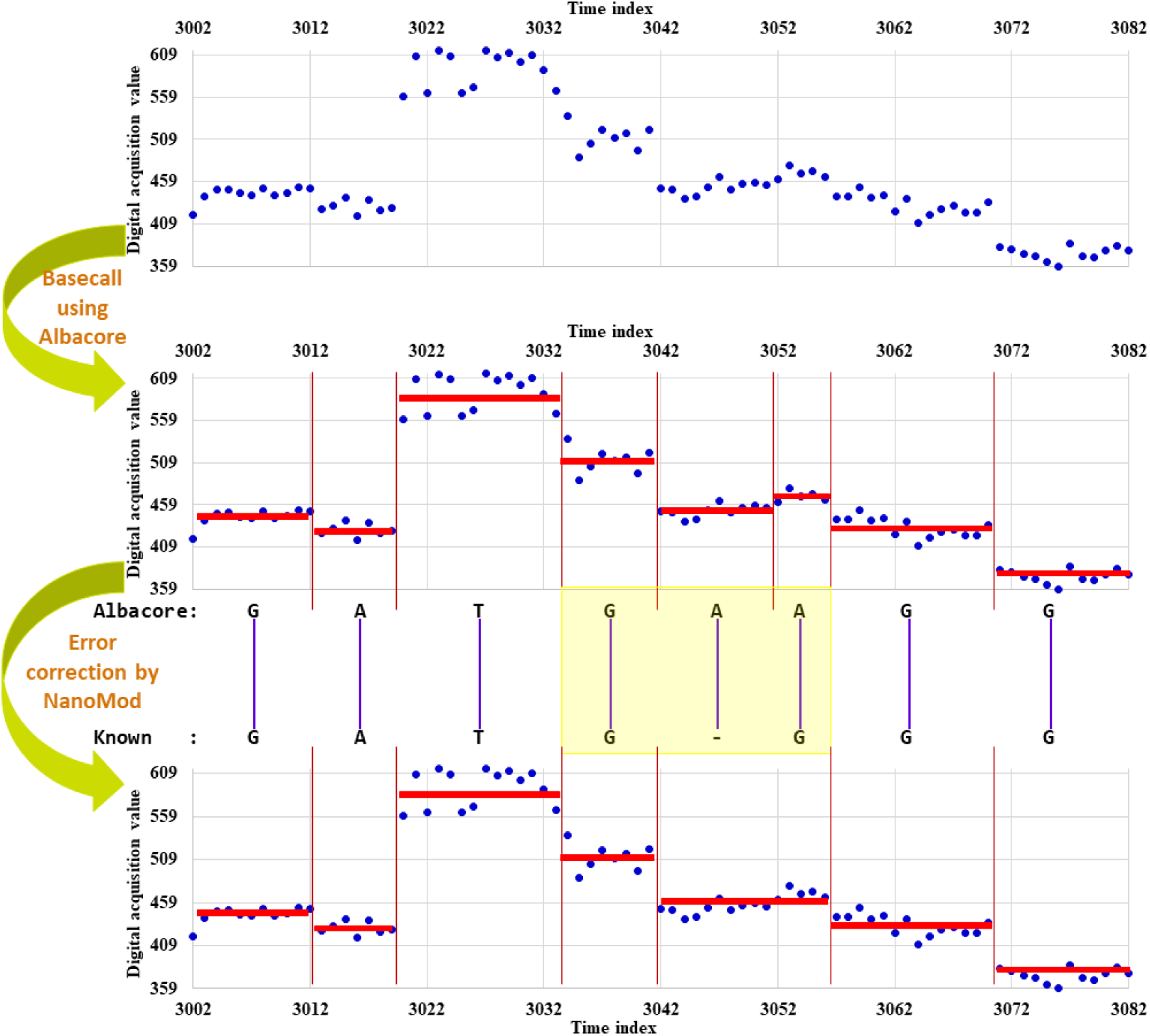
*An example of the insertion correction procedure in NanoMod*. *The region shadowed in yellow shows the insertion base together with one upstream and one downstream neighbors. For other notations, see Figure 2.*

After that, raw signals in a long read are normalized using the median subtraction and the standardization by averaged difference, and the normalized signal was limited between −5 to 5. Normalized signal information of each position in a long read subsequently anchors a position in the known reference sequence. This process is similar to what is described in nanoraw[2].

### Signal summarization for positions in the known sequence

Based on the corrected alignment of a long read with the known sequence, the normalized signal of a position in a long read can be assigned to the corresponding aligned position in the known sequence. Given two groups of aligned long reads, each position in the know sequence will have two groups of normalized signals, one from reads of the sample with modifications and the other from the matched non-modified sample.

Sometimes, a position may have a much smaller number of associated reads in one sample versus the other sample, possibly due to random fluctuation of coverage or due to other issues (for example, PCR amplification biases). Thus, those positions with limited data on signals in either group are filtered and excluded from the downstream analysis, based on user-specified criteria.

### Detection of modifications

Assuming that signals of a base for a position in a known sequence are generated from a specific but unknown distribution with some noises. The signals for a position of the known sequence in the two groups would be highly similar to each other if the position and its closest neighbors are not modified. However, if a position contains a modified base, the signals of the two groups for the position and/or its neighbors would be different, in term of mean, standard deviation or shape. In other words, a position has high probability to have a modified base if the signals between the two groups for the position or its neighbors are statistically different.

In NanoMod, Kolmogorov-Smirnov test is used for this purpose, since our purpose is to detect *de novo* modifications and since the actual distribution of signal intensity is not known *a priori*. Additionally, our experience and manual examination showed that the distribution of signal intensities at a modified position (or neighbors of a modified position) can be of various different shapes, such as increased/decreased mean, increased variance, a change from unimodal to bimodal distribution, etc. Kolmogorov-Smirnov test [22] is one of the most useful nonparametric test methods to quantify the distance between empirical distribution functions of two groups of samples. It is sensitive to the differences in both the locations and shapes of the two distribution functions. The Kolmogorov-Smirnov statistic *D*_*m,n*_ is defined below.

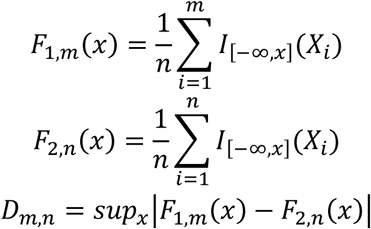

Where *X*_*i*_ is a signal, and *I*_[−∞,*x*]_(*X*_*i*_) is 1 if *X*_*i*_ ≤ *x* and 0 otherwise. *F*_1,*m*_(*x*) is for a group of *m* modified reads, and *F*_2,*n*_(*x*) is for a group of *n* non-modified reads. *sup* is a supremum function giving the least upper bound, that is, the least difference which is not less than all differences between the two *F*(*x*)s. P-values of the Kolmogorov-Smirnov test indicate the probability of the base at a position to be modified: the smaller p-value is, the more likely the base is modified.

### The combination of neighbor p-values

Measured signals in Nanopore data are usually for k-mers, that is, a modification of a base at a specific position may affect the signals of its neighbors. Therefore, p-values of neighboring positions may also suggest the presence of modifications. To take into account the neighborhood effect, p-values within *k* closest positions of a given position can be used to generate a combined p-values. *k* could be specified by users and by default *k =* 2. Weighted Stouffer’s method is used for this purpose, so that the center position has higher weights, and the further neighbors, the lesser the weights. The weighted Stouffer statistic for *k* + 1 consecutive positions (*k* closest positions plus the center position) is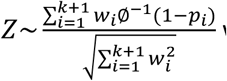 where *p*_*i*_ is the probability of a position with a weight of *w*_*i*_ and ∅*−*^1^(1 − *p*_*i*_) returns a Z score of *p*_*i*_ with a standard normal cumulative distribution function.

When a position has extremely small p-value, its neighboring positions tend to also have very small p-values, and these positions will rank very high among all positions. Therefore, the rank for a position gives redundant information on whether a neighborhood region has a modification. We thus used neighborhood-based ranking. In neighborhood-based ranking, if a position has a higher rank, its neighbor positions (within 1 or 2 base window size for both left and right sides) with lower rank are not considered.

### Simulation of nanopore long-read data

To evaluate how NanoMod works on modifications with different properties, we generated several simulation datasets where samples have multiple types of modifications. In the simulation, we assumed that we had a sequence and each 5-mer produces signals according to a normal distribution of the mean E_*k*_ and the standard deviation Δ_*k*_ plus some random noises, then a basic simulation process for a given sequence can be described as below:

1. Generate n signals for each 5-mer in the given sequence, and sequentially merge all signals together for the given sequence. n is a random number which varies from 5 to 15.
2. Repeat Step 1 for 100 times, and treat them as raw reads of a non-modified sample.
3. Sample h positions in the given sequence, and assume that those bases are modified.
4. For each position h_*i*_ with simulated modifications and its neighborhood position h_*j*_, ‖*j - i‖* ≤ 2, the mean was increased by 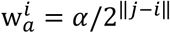, and the standard deviation was increased by 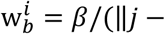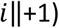 If a position is adjacent to two modifications, *h*_*u*_ and h_*v*_, its 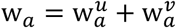 and 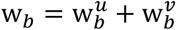 otherwise if a position is only close to a modifications h_*i*_, 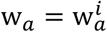 and 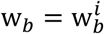 In this study, α was set to 0.2, while β was set to 1.
5. For those positions with modifications or are adjacent to the modified bases, generate m signals according to a normal distribution of the mean E_*k*_ * (1 + w_*a*_) and the standard deviation Δ_*k*_ * (1 + *w*_*b*_) plus some random noises. Here E_*k*_ and Δ_*k*_ are the mean and standard deviation of the corresponding non-modified 5-mer, and m was a random number, which varies from 5 to 15.
6. For other positions without modified bases and are not in the vicinity of modified bases, generate m signals as what has been done in Step 1.
7. Repeat Steps 4, 5 and 6 for 100 times, and treat them as reads of a modified sample.
8. Run NanoMod on two groups of reads.
9. Repeat Steps 1 to 8 for 100 times so that 100 pairs of datasets were used to evaluate NanoMod.

To simulate modifications with different properties, we generated several types of simulation data sets below:

i. ‘MeanDif’ simulation: The modification of a base only affects signal mean of the 5-mer centered at that base, *i.e., w*_*a*_ > 0. Signal standard deviation of the 5-mer has no change (*w*_*b*_ *=* 0) and no neighborhood effect (*w*_*a*_ *=* 0 *and w*_*b*_ *=* 0 for non-modified bases).
ii. “STDDif” simulation: The modification of a base only affects signal standard deviation of the 5-mer centered at that base, *i.e., w*_*b*_ > 0. Signal mean of the 5-mer has no change (*w*_*a*_ *=* 0) and no neighborhood effect (*w*_*a*_ *=* 0 *and w*_*b*_ *=* 0 for non-modified bases).
iii. “Mean_STDDif” simulation: The modification of a base affects both signal mean and standard deviation of the 5-mer centered at that base, *i.e., w*_*a*_ > 0 *and w*_*b*_ > 0, but no neighborhood effect (*w*_*a*_ *=* 0 *and w*_*b*_ *=* 0 for non-modified bases).
iv. “Mean_STDDif_NE” simulation: The modification of a base affects both signal mean and the standard deviation of the 5-mer centered at that base, *i.e., w*_*a*_ > 0 *and w*_*b*_ > 0, and also adjacent neighbors, *i.e.* > 0 *and w*_*b*_ > 0 for adjacent non-modified 5-mer of the modified bases.

### A Nanopore long-read sequencing data set on *E. coli*

A publicly available Nanopore long-read sequencing data of *E. coli* [3] was also used to evaluate NanoMod. This dataset contains two groups of samples, one was generated from PCR product where DNA modifications are not expected to be present, and the other was from PCR product then subject to enzymatic methylation with the M.SssI methyltransferase where nearly all of cytosines in a CpG context were converted to 5-mC. These dataset was downloaded from the European Nucleotide Archive under accession number PRJEB13021 [3]. On this data set, the known *E. coli* sample has ∼4.64Mb nucleotides and ∼693,586 CpG sites.

### Measurement for performance evaluation

To measure the performance of ranking modified bases at the top among all bases, we used the percentiles of 0.1%, 0.25%, 0.5%, 1%, 2%, 3%, 4% and 5% to split the ranking into 9 categories for simulation data. Then, at each percentile, we calculated precision (*i.e.*, the number of correctly identified modifications divided by the number of modification predictions at a percentile) and recall (*i.e.*, the number of correctly identified modifications divided by the number of modifications) for correctly detecting the known modifications, and generated precision-recall plot. On the *E. coli* data set [3], we used the percentiles of 0.1%, 0.25%, 0.5%, 1%, 2%, 3%, 4%, 5%, 10%, 15%, 20%, 25% and 30% to split the ranking, because there are many methylated CpG sites.

## Results

NanoMod was designed to detect candidate positions with DNA modifications using raw electric signals generated from Nanopore long-read sequencing techniques. Briefly, two groups of reads, one containing modified bases and the other without modified bases, were used as input of NanoMod, and were then subject to basecalling, error correction and signal annotation of positions in a known sequence. After that, we rank all positions for the presence of potential DNA modifications. NanoMod was evaluated on simulation data where raw signals were simulated according to the mean and standard deviation of 5-mer and on a publicly available methylation data. The results are described in detail below.

### Evaluation on simulation data with multiple DNA modifications in a sequence

We simulated 200 reads of a 6184-bp sequence (100 with and 100 without modification on specific positions) on a sample based on the means and standard deviations of observed 5-mers (1,024 distributions) from large-scale Nanopore sequencing experiments. Each of the simulated modification reads has 60 modifications randomly dispersed across the whole sequence. DNA modifications with different properties were simulations as described in the Method section. Each type of modification was also generated 100 times. The performance of NanoMod was evaluated using precision and recall, as the percentile of the rank which ranges from 0.1%, 0.25%, to 0.5%, 1%, 2%, 3%, 4% and then to 5%. Typically, when the percentile value increases, the recall would increase. We compared the performance of Mann–Whitney U test, Student’s T test and Kolmogorov-Smirnov test on signals of single bases, and also two combined statistics methods including Stouffer’s method and Fisher’s method. The results were shown in **Figure 4**, where precision and recall were the averaged values on 100 simulation data sets.

**Figure 4.**
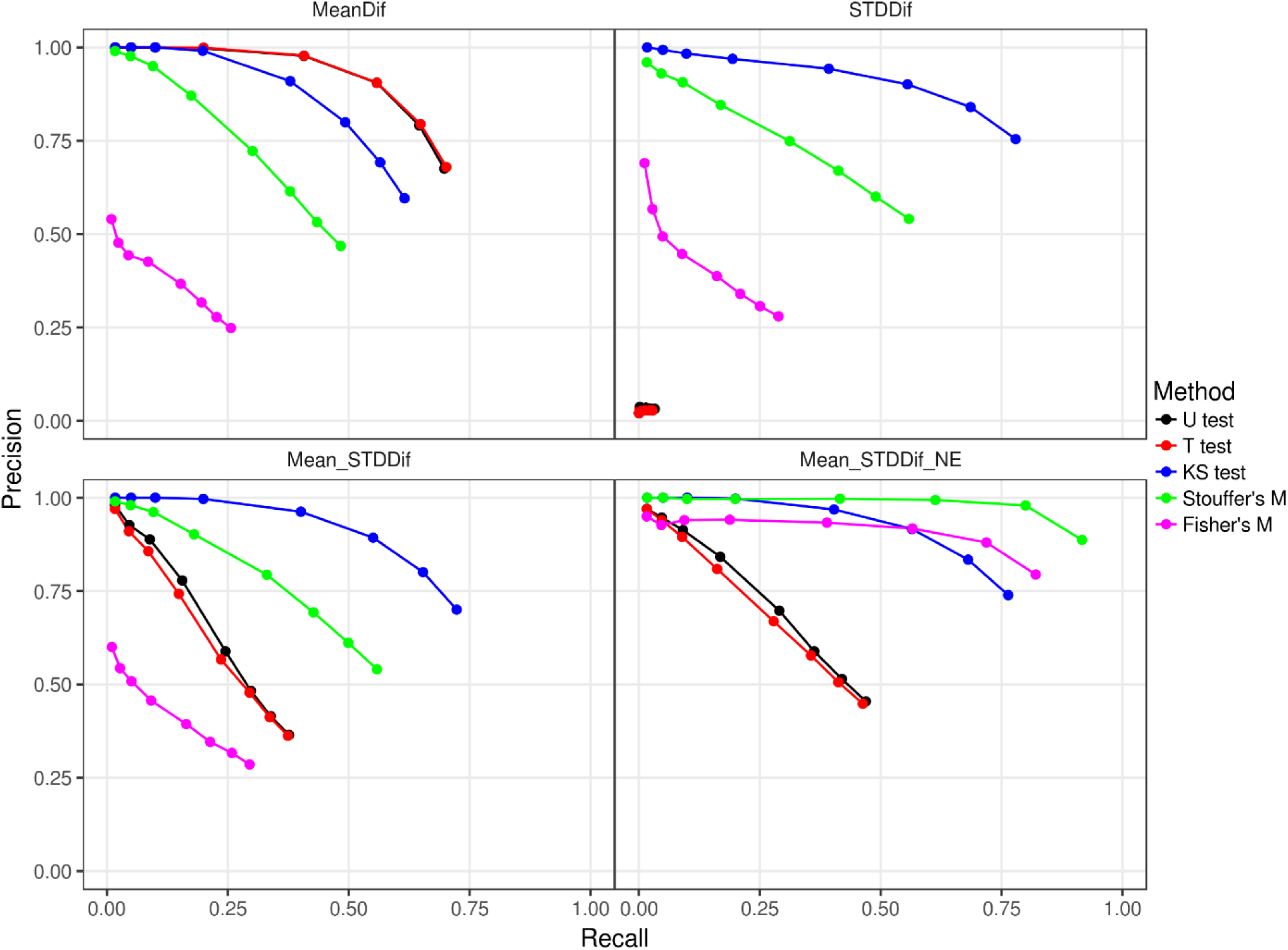
The average precision and recall rates of NanoMod on 100 simulation data sets. “U test”:Mann–Whitney U test, “T test”:Student’s T test, “KS test”:Kolmogorov-Smirnov test, “Stouffer’s M”: Stouffer’s method and “Fisher’s M”:Fisher’s method. “MeanDif”: modified bases only have the mean difference for signals from non-modified bases, “STDDif”: modified bases only have the difference of standard deviation for signals from non-modified bases, “Mean_STDDif”: modified bases have the difference of the mean and standard deviation for signals from non-modified bases, and “Mean_STDDif_NE”: the simulation of “Mean_STDDif” plus neighborhood effect. Precision was calculated using the number of correctly identified modifications divided by the number of modification predictions at a percentile (i.e., 0.1%, 0.25%, 0.5%, 1%, 2%, 3%, 4% and 5%), and recall was calculated using the number of correctly identified modifications divided by the number of modifications.

We found that on the MeanDif simulation (see the Method section), Mann–Whitney U test and Student’s T test worked better than Kolmogorov-Smirnov test, because the first two statistics methods were more powerful to detect the change of the mean of 5-mers, yet signal mean of modified bases were simulated on the MeanDif simulation. However, for the other three simulations (the STDDif, Mean_STDDif, and Mean_STDDif_NE simulations) where differences of standard deviation of 5-mers were considered, Kolmogorov-Smirnov test significantly outperformed Mann–Whitney U test and Student’s T test. In particular, when only differences of standard deviation of modified 5-mers were simulated, Mann–Whitney U test and Student’s T test had no predictive power, as expected (refer to STDDif in **Figure 4**). Since it is unusual for DNA modifications to change only mean or variance of signal intensity values, Kolmogorov-Smirnov test is used by default in NanoMod for capturing all types of alterations in signals.

When the two combined statistics methods were used, the performance was worse than Kolmogorov-Smirnov test for the MeanDif, STDDif and Mean_STDDif simulations. This is because no neighborhood effects were considered in these types of simulations. When neighborhood effects were simulated in Mean_STDDif_NE, both Stouffer’s Method and Fisher’s method improved the detection of DNA modification. In particular, Stouffer’s method performed much better than Fisher’s method, and 75% modifications detected by Stouffer’s method were ranked at top. This suggested that Stouffer’s method is preferred over single-base Kolmogorov-Smirnov test when neighborhood effects are present.

### Evaluation on *E. coli* methylation data

To test the usefulness of NanoMod on synthetically introduced modifications rather than simulated data, we also evaluated NanoMod on a publicly available Nanopore long-read sequencing data on *E. coli* [3] where CpG sites were almost all methylated by the M.SssI methyltransferase. Given a rank list of detected modifications, we calculated precision and recall at each splitting percentile value for evaluating five statistical methods and nanoraw. The results were shown in **Figure 5**.

**Figure 5.**
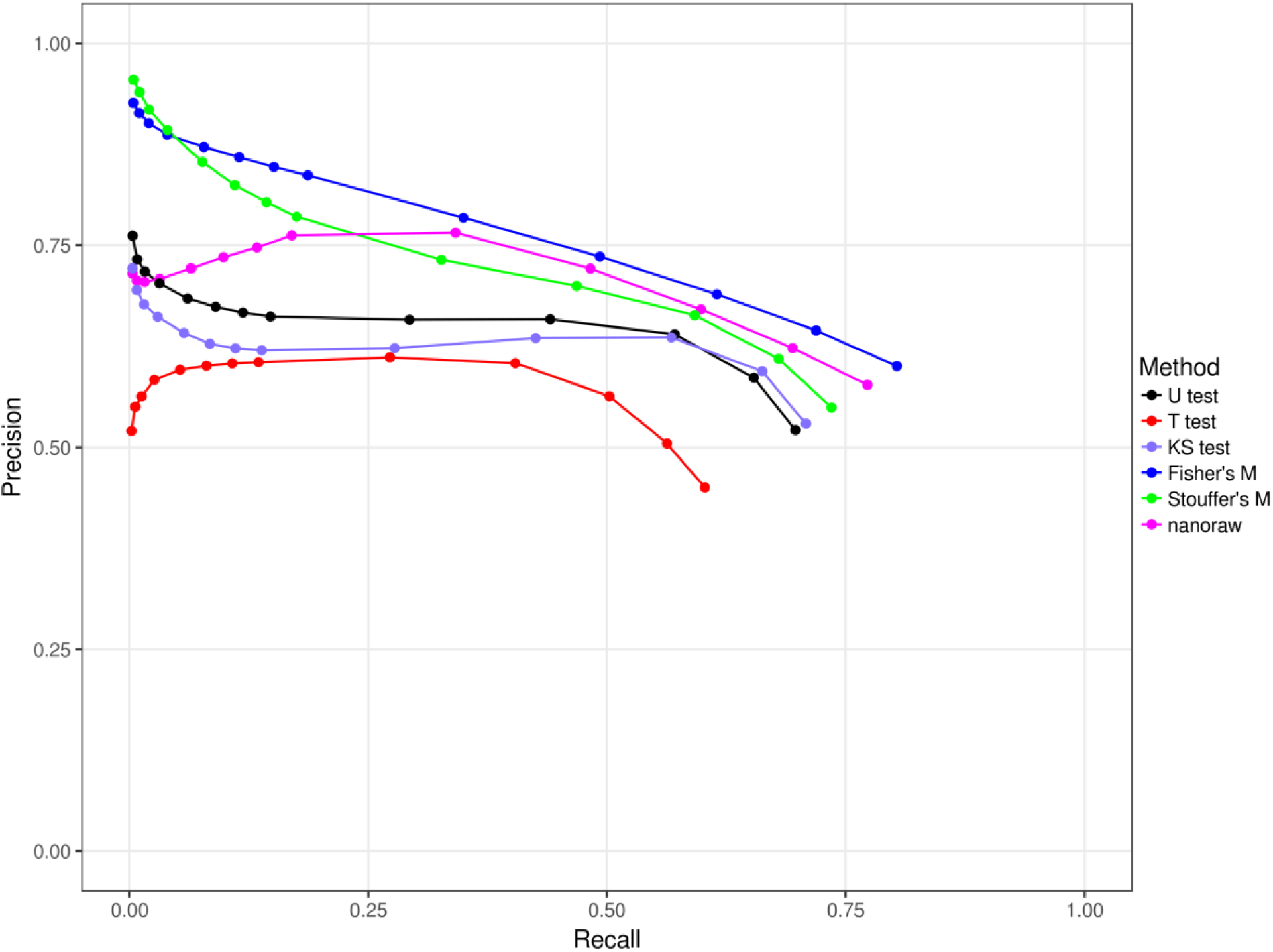
The performance of modification detection using nanoraw and 5 statistics methods implemented in NanoMod. “U test”: Mann–Whitney U test, “T test”: Student’s T test, “KS test”: Kolmogorov-Smirnov test, “Stouffer’s M”: Stouffer’s method and “Fisher’s M”: Fisher’s method. The percentile definition was in the method section. Precision was calculated using the number of correctly identified modifications divided by the number of modification predictions at a percentile (i.e., 0.1%, 0.25%, 0.5%, 1%, 2%, 3%, 4%, 5% and then to 30%), and recall was calculated using the number of correctly identified modifications divided by the number of modifications.

We found that the combined statistical testing methods achieved better performance than the various methods on single bases, indicating strong neighborhood effects caused by the methylation. In particular, Fisher’s method and Stouffer’s method in NanoMod outperformed nanoraw especially to detect methylations at the top rank (the smaller recall in **Figure 5**), where the precision of nanoraw was about 0.70 while the precision of Nanomod was more than 0.9. (We note that nanoraw itself incorporated Fisher’s method to combine p-values.) Therefore, NanoMod significantly improved the performance to detect modified DNA bases.

In **Figure 5**, the better performance of Fisher’s method is due to the fact that the *E. coli* sequence has many modified regions containing multiple CpG sites together. Note that the original data set was generated on DNA molecules treated with the M.SssI methyltransferase, where nearly all of cytosines in a CpG context were converted to 5-mC, yet in practice *de novo* modifications in living organisms should occur only in a very minor fraction of bases. Indeed, upon further examination, among about half of CpG sites, we can found another CpG sites within 5 bps. In such scenarios, Stouffer’s method cannot work as well as Fisher’s method, because Stouffer’s method give less weights to neighbors of a methylation site.

### Example plots of top ranked CpG sites

We used the top three ranked CpG sites detected by NanoMod in the methylation data of *E. coli* to demonstrate details of how NanoMod identified signal difference at methylated CpG sites. The results were shown in **Figure 6**.

**Figure 6.**
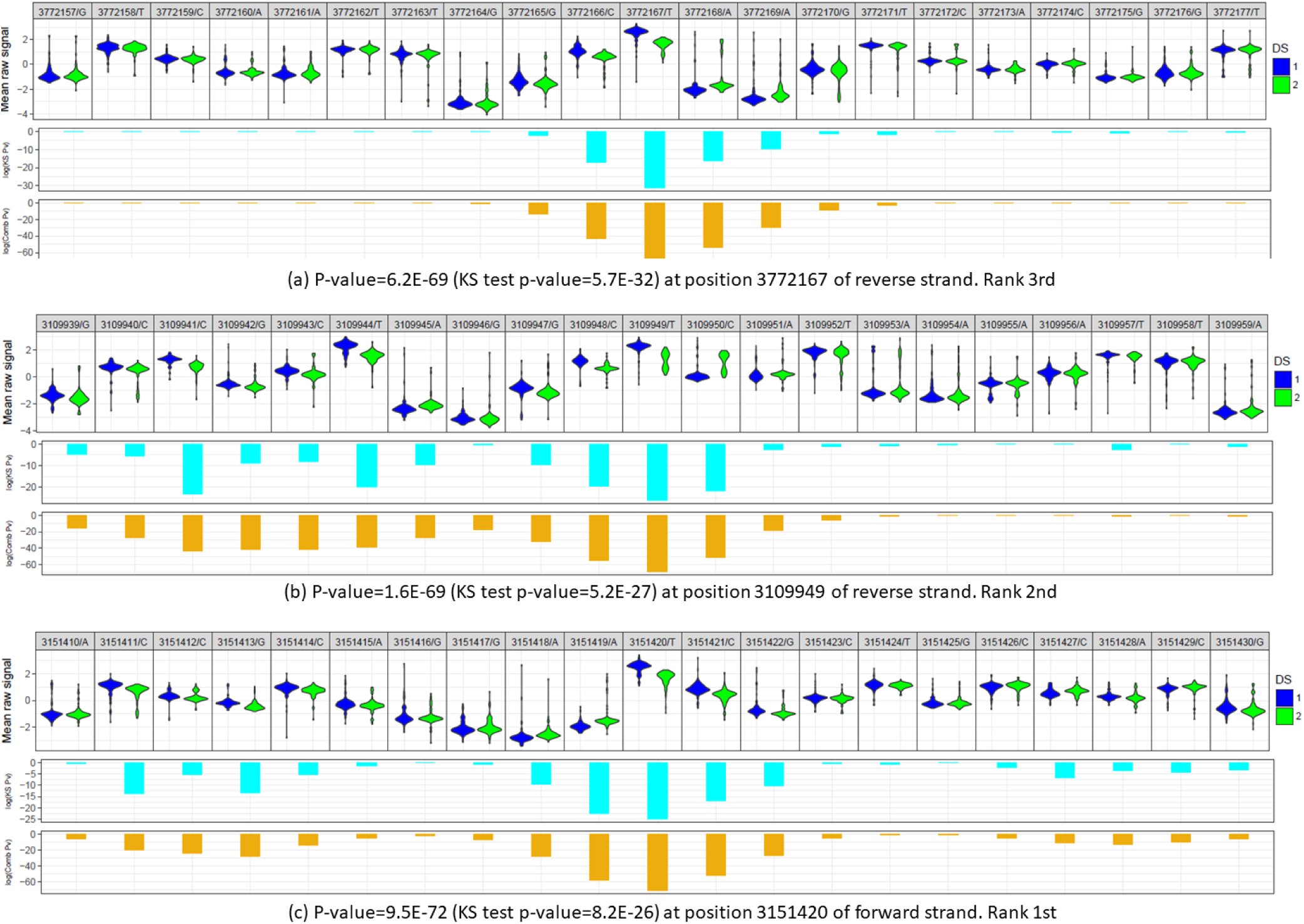
*Analysis of DNA methylation in E. coli using NanoMod*. *Data of the top three ranked modifications are shown. ‘p-value/Comb Pv’ is the combined p-values calculated using the Stouffer’s method, ‘KS test/KS Pv’ is the p-value calculated using Kolmogorov-Smirnov test. ‘DS 1’ represents the non-methylated sample, and ‘DS 2’ represents the methylated sample. In each panel, the first line is the position of the base in the reference genome followed by the base in reads. The position is based on the reference genome. For reverse strand, the 3’ to 5’ of reads is from right to left while for forward strand of reads is from left to right.*

As can be seen from **Figure 6** (a) and (b), NanoMod generated the smallest p-values of the position and its closest neighbors at the center of the plot. In contrast, those positions, which were far from the CpG sites (the both left and right sides of **Figure 6** (a) and the right side of **Figure 6** (b)), had much larger p-values using either Kolmogorov-Smirnov test or the Stouffer’s method. This observation demonstrated that methylated bases changed the Nanopore signals. In the left side of **Figure 6** (b) and **Figure 6** (c), the methylated CpG site in the center had smaller p-values. Meanwhile, the left side of **Figure 6** (b) and the left and right sides of **Figure 6** (c) also had p-values which were smaller than the both left and right sides of **Figure 6** (a) and the right side of **Figure 6** (b). This is because there are two additional CpG sites on the right side of **Figure 6** (b) and two additional CpG sites on either side of **Figure 6** (c). These observations clearly demonstrated that NanoMod captured statistically significant signals of modified bases between modified reads and non-modified bases.

## Discussion

The advent of Nanopore long-read sequencing technique provides valuable opportunities to detect DNA modifications directly from signal intensity data at a large scale and at low costs. Although several existing tools were developed for the detection of DNA modifications using Nanopore long-read data, they either need large training data [3, 20, 21] or require further tweaking of algorithms to improve the detection of modifications [2]. In this study, we proposed NanoMod to identify DNA modifications using raw signals of reads generated by the Nanopore long-read sequencing technique. NanoMod does not require any training data so it can detect DNA modifications *de novo*. NanoMod achieved improved performance when it was evaluated on simulation data with modifications of different properties and on an *E. coli* data set with 5mC (5-methylcytosine) modifications.

Compared with existing methods [3, 20, 21], one limitation of NanoMod is that it is designed to detect DNA modifications *de novo* and hence cannot predict the specific type of modification (such as whether a modification is 4mA or 5mC). However, given large-scale training data sets, it is possible to generate prior models in NanoMod to detect specific type of modifications. For example, assume that a specific modification on a base will impact the signals on a 7-bp window around the base, then from a large training data set, we can generate the signal distributions for all 7-mers surrounding the specific modification as the prior model. Long reads on a new sample can be compared to the prior model as well as a control model (from samples without modifications) to detect whether the specific type of modification exist in each position in the new sample.

Both nanoraw [2] and NanoMod require two groups of reads, one from a sample with modifications and the other from a matched control sample known to contain no modifications. They share one critical component, which is the error correction procedure from the alignment. After aligning long reads to a known sequence, signals of bases in long reads can be used to annotate the corresponding mapped positions. However, Nanopore long reads have high insertion/deletion errors especially when the bases are called from segmentation events (the latest version of albacore no longer use this procedure, partly due to the inherent error in the segmentation process). Thus, the indel correction step is crucial to rescue signal annotations for many positions. In some specific cases, the reference sequence used in NanoMod may contain true indels themselves (for example, when using *E. coli* reference genome sequence in the analysis for an *E. coli* strain with real indels at specific positions); in these scenarios, it may be necessary to generate confidence sets of indel calls on the sample first, then use a modified version of reference sequence (by incorporating highly confident indels) in the error correction procedure in NanoMod.

Integrated statistical testing is another critical component of NanoMod, because different types of modifications can result in different types of alterations on the signal distributions. Modifications might (i) only change signal mean of a modified base, or (ii) only change variance of the signals for a modified base, or (iii) result in the change of both signal mean and standard deviation, or (iv) change the overall shape of distribution such as from unimodal to bimodal, or (v) affect several adjacent neighborhood positions, and so on. Different statistical tests may have different power to detect modifications based on the property of the modifications. For example, Student’s T test and Mann–Whitney U test outperformed Kolmogorov-Smirnov test for the first category of modifications, but had no prediction power for the second category of modifications and had limited power for the other categories of modifications (see **Figure 4**). Kolmogorov-Smirnov test outperformed other methods for the second and third categories of modifications, and achieved worse performance than the combined statistical testing for the fifth category of modifications. In short, Kolmogorov-Smirnov test on single bases and the combined Stouffer’s method on multiple bases are better choice for modification detections than Mann–Whitney U test used in nanoraw [2] when data set is large enough. This comparison was supported by additional preliminary studies (data not shown) where NanoMod could achieve much better prediction performance than nanoraw for multiple different types of modifications that we introduced synthetically into DNA molecules.

More accurate detection of DNA modifications, especially synthetically introduced modifications, has many downstream applications. For example, accurate detection of DNA modifications can facilitate studies on the role of epigenetic modifications [24] in different human diseases such as cancer, and help identity candidate genes where epigenetic switch is important for disease progression. Similarly, incorporation of modified DNA bases during DNA synthesis can be used to track patterns of DNA replication [25, 26], so accurate detection of *de novo* DNA modifications on newly synthesized DNA strands enables genome-wide studies on DNA replication timing and patterns.

## Conclusion

We have developed a new computational tool, NanoMod, for the detection of DNA modifications using Nanopore long-read sequencing data. We evaluated NanoMod on simulation data with different types of modifications and also on a methylation data of *E. coli*. Our results suggested that NanoMod achieved better performance than other existing tools in detecting modifications without training data. Therefore, NanoMod will greatly facilitate functional genomics experiments for single base resolution mapping of modified nucleotides in the genome.

## Declarations

### Authors’ contributions

QL and KW designed the study. QL implemented the tool and performed the analysis. DCG and DME involved in the improvement of the tool and analysis. QL and KW drafted the manuscript. All authors read, revised and approved the manuscript.

## Acknowledgements

We thank the Wang lab members for insightful comments on the NanoMod algorithm and implementation, and we thank the developers of nanoraw for providing valuable help and support on the software tool. We thank Simpson *et al* for making the *E. coli* data publicly available for testing our software tools.

## Declaration

This study was supported by NIH grant HG006465 (K.W.).

